# Coevolution in Small Heat Shock Protein 1 (HSPB1) is Promoted by Interactions between the Alpha-Crystallin Domain and the Disordered Regions

**DOI:** 10.1101/2024.12.03.626587

**Authors:** Vanesa Racigh, Luciana Rodriguez Sawicki, Facundo Nicolas Eric Bravo, Maria Silvina Fornasari

## Abstract

Human HSPB1, a member of the small heat shock protein (sHSP) family, functions as an ATP-independent molecular chaperone crucial for protein quality control and is implicated in several pathologies, including Charcot-Marie-Tooth neuropathy. This study investigates the coevolution of the disordered N-terminal and C-terminal regions (NTR and CTR) with the structured Alpha-Crystallin domain (ACD) of human HSPB1, focusing on interactions that regulate its chaperone activity.

Using a manually curated dataset of HSPB1 orthologs, the composition of critical motifs within the NTR (_6_VPFSLL_11_) and CTR (_179_ITIPV_183_) that interact with the ACD was analyzed and evolutionary rates per site for the human HSPB1 sequence were estimated. Additionally, structural modeling with AlphaFold 2 was employed to assess the prevalence of these contacts in human HSPB1 models.

The results reveal that while the disordered regions globally evolve faster than the structured ACD, specific residues within the _6_VPFSLL_11_ and _179_ITIPV_183_ motifs exhibit reduced evolutionary rates, reflecting evolutionary constraints imposed by the conservation of the protein’s function. Structural modeling further indicates that coevolutionary-like information about the interaction between the _6_VPFSLL_11_ motif and the ACD is encoded in the multiple sequence alignment used by Alphafold 2.

Altogether, these findings suggest that the disordered regions and the ACD of human HSPB1 likely coevolved, preserving interactions crucial for its chaperone activity self-regulation. This evolutionary mechanism may also be extended to other sHSP featuring interacting motifs in the NTR, CTR, or both, and provides a framework to elucidate why pathogenic variants occurring in regions involved in these contacts contribute to disease.

## Introduction

Small heat shock proteins (sHSPs) are ATP-independent molecular chaperones that act as the first line of defense in the cellular chaperone network [1]. The human genome encodes ten sHSPs (HSPB1 to HSPB10), which can be ubiquitously expressed or exhibit tissue-specific expression patterns and become upregulated under conditions of cellular stress. Their primary role is to interact with misfolded clients to prevent their aggregation. Variants of these proteins with altered chaperone activity have been linked to several diseases, including Parkinson’s, Alzheimer’s, and various neuropathies [2].

Structurally, sHSPs consist of a conserved Alpha-Crystallin domain (ACD), flanked by disordered N-terminal (NTR) and C-terminal (CTR) regions [3]. sHSPs can form dynamic homo- and hetero-oligomers of varying sizes, which are stabilized by extensive contacts between the disordered regions and the ACD [1,4,5]. Under stress conditions, specific phosphorylation sites in the NTR trigger the disassembly of oligomers into smaller species, freeing the NTR from oligomeric contacts and enhancing its interaction with exposed hydrophobic regions of misfolded proteins [6,7]. Experimental studies indicate that the dimeric form of sHSPs exhibits the highest chaperone activity, making dimer constructs the standard unit for assessing chaperone function [8]. Each ACD dimer presents three grooves: a central groove at the dimer interface and two lateral grooves, one on each subunit. These grooves serve as interaction sites for specific motifs within the disordered regions, such as the I/V-X-I/V motif in the CTR and the conserved SRLFDQXFG motif in the NTR, which are implicated in oligomerization [3,9,10]. In higher-order assemblies, the CTR often functions as a non-covalent “cross-linker” between dimers by binding the I/V-X-I/V motif into the lateral groove of a neighboring dimer [5,11,12]. This interaction is crucial for oligomer formation.

Recent work by Clouser et al. provides a detailed experimental study describing the interactions between the NTR and the ACD of human HSPB1 [13]. They found, following the canonical human HSPB1 sequence and sequence numbering, that the _6_VPFSLL_11_ motif located at the distal part of the NTR interacts with the lateral grooves in a dimeric construct (Figure 1). Interestingly, it has been shown that abolishing this interaction enhances in vitro human HSPB1 chaperone activity towards its natural client tau [14]. Additionally, the crystal structure of the oligomeric form of human HSPB1 reveals that an overlapping or “extended” I/V-X-I/V motif (_179_ITIPV_183_) within the CTR also interacts with the ACD’s lateral grooves, highlighting the importance of these contacts [5]. Moreover, the substitutions P7R, P7S, P182A, and P182L, that occur at positions within these motifs, have been associated with Charcot-Marie-Tooth disease [15–20].

**Figure 1.**
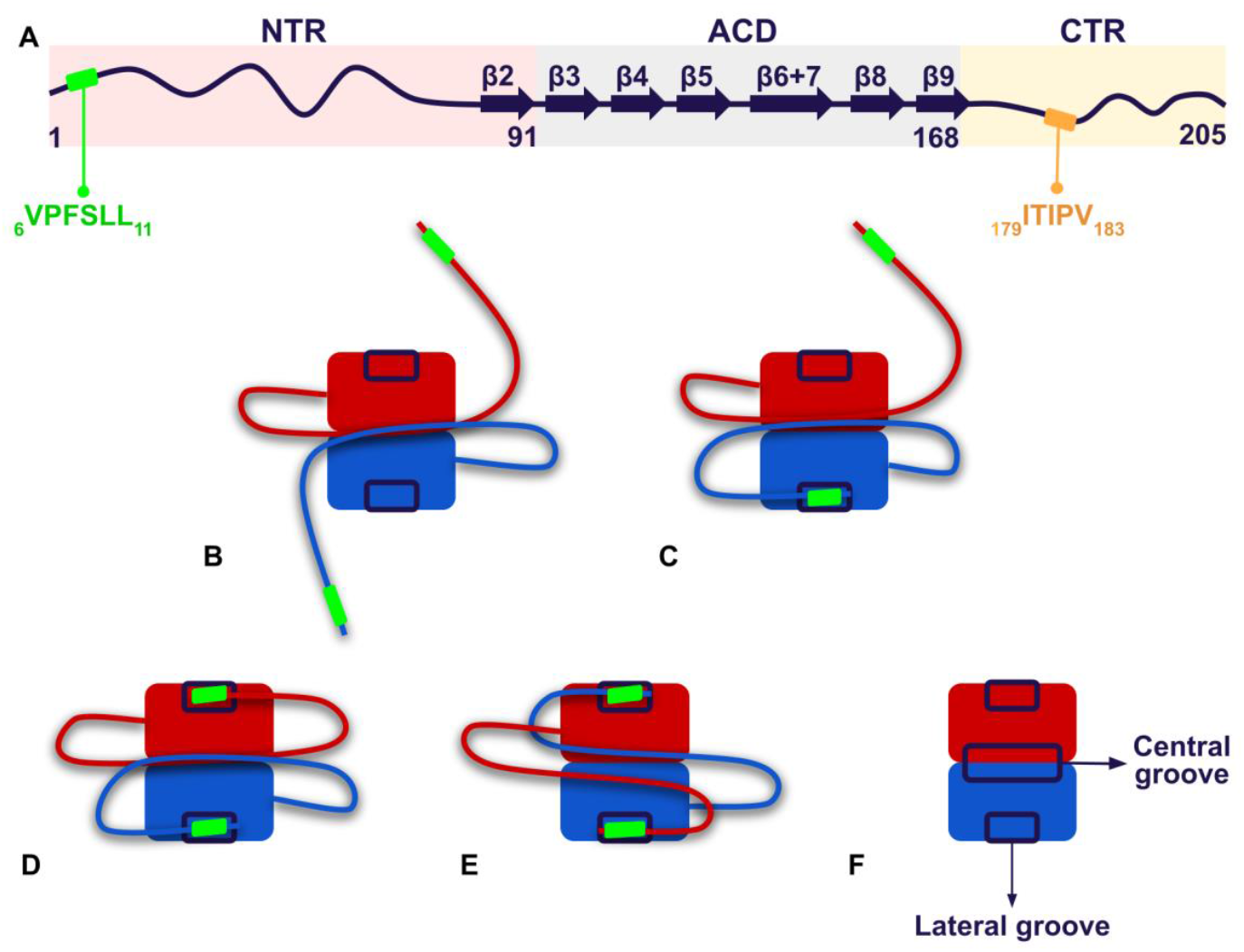
Schematic representation of the human HSPB1 structure and the interactions between the distal end of the NTR and the lateral grooves of the ACD via the _6_VPFSLL_11_ motif in the dimer, as described by Clouser et al. [13]. **A**. The primary and secondary structure of human HSPB1 is shown, indicating NTR, ACD, and CTR regions, as well as the _6_VPFSLL_11_ and _179_ITIPV_183_ motifs. **B-E**. The figures illustrate five possible interaction modes of the NTR’s distal region within the human HSPB1 dimer: **B**. both NTRs are free in solution; **C**. the NTR of one chain interacts with the lateral groove of the same chain while the NTR of the other subunit remains free; **D**. both NTRs are bound to the lateral grooves of their respective chain; and **E**. the NTR of each chain is in contact with the lateral groove of the opposite chain. In all modes, the CTR is not represented, while the global conformation of the NTR is depicted schematically, indicating the _6_VPFSLL_11_ as a green rectangle. **F**. ACD dimer highlighting the lateral grooves that interact with the distal end of the NTR, and the central groove, which contacts other NTR regions, including the conserved motif (_26_SRLFDQXFG_34_ in human HSPB1). Baughman et al. found that chaperone activity increases when the distal NTR is released from the lateral grooves of the ACD (activated state, as in **B** and **C**), suggesting that this interaction restricts the conformations of the NTR required to effectively perform chaperone activity [14].

These findings underscore the crucial role of the interactions between the _6_VPFSLL_11_ and _179_ITIPV_183_ motifs, located in the disordered regions, and the ACD in the self-regulation of human HSPB1 chaperone activity. Both oligomer formation and the reversible interactions between the NTR and the ACD at the dimeric level are fundamental to this regulatory process. Given the current evolutionary hypothesis suggesting that the disordered regions of sHSPs evolved independently and at a faster rate than the ACD [21], it is relevant to explore how these contacts have shaped the evolution of human HSPB1. To investigate the evolutionary importance of these interactions, we used a manually curated dataset of HSPB1 orthologs to estimate the global evolutionary rates for the NTR, ACD, and CTR, and to calculate site-specific evolutionary rates by taking the human HSPB1 sequence as a reference. Furthermore, we derived evolutionary insights from models generated with AlphaFold 2 (AF2) [22]. Our results suggest that the ACD, NTR, and CTR of human HSPB1 did not evolve independently; rather, they must have coevolved to maintain the critical interactions necessary for regulating its chaperone activity.

## Methods

### HSPB1 orthologs dataset creation

Homologous protein sequences were initially recruited from the UniprotKB database [23] with Uniprot BLASTp [24], using the canonical human HSPB1 (UniProt ID P04792) as a query. To ensure a comprehensive dataset, additional sequences were sourced by filtering according to specific taxonomic groups. The recruitment process followed the phylogenetic tree provided by Hedges et al. [25], focusing on major kingdoms: Animalia, Plantae, Eubacteria, Fungi, and Protista. When the maximum number of sequences was reached for a kingdom, further recruitment was conducted within subgroups of that kingdom to capture broader diversity. After recruitment, an initial filtering step was applied to remove sequences that were partial, hypothetical, or contained indeterminate residues. Duplicated sequences were also eliminated to avoid redundancy. The remaining sequences were aligned and then filtered based on percentage identity and sequence coverage using an in-house program that implements Biopython [26]. A threshold of 30% identity and 40% coverage was applied. From the resulting dataset, all HSPB1 sequences were manually curated to compile a set of orthologs HSPB1 sequences for subsequent analysis. The importance of working with proteins coded by orthologs genes is related to studying the evolutionary history of the product of a single gene, ensuring that these are the same protein in different organisms and, consequently, they perform the same function.

### Dataset characterization

Two subsets were generated from the orthologs dataset, one for vertebrates and another for invertebrates. The three sequence datasets (the complete set, the vertebrate subset, and the invertebrate subset) were aligned using Clustal Omega within UGENE [27]. From these alignments, the amino acid composition and the observed substitutions of the _6_VPFSLL_11_ motif within the NTR and the _179_ITIPV_183_ (I/V-X-I/V variant) in the CTR were analyzed. Additionally, the amino acid composition of the β4 and β8 strands, which frame the lateral grooves (positions _109_LTVKT_113_ and _153_VSSSL_157_, respectively), was examined.

### Structural and evolutionary analysis

Evolutionary distances within the NTR, CTR, and ACD datasets were calculated using the protdist program from the PHYLIP package with default parameters [28], following the method described by Kriehuber et al. To visualize the distribution of these evolutionary distances, we applied Kernel density estimation to the flattened distance matrices. Differences in evolutionary rates between these regions were statistically assessed using the Kolmogorov-Smirnov (KS) test implemented in Python. The evolutionary rate per site was calculated using the Rate4Site program with default parameters [29]. The analysis was conducted on the protein sequences of HSPB1 orthologs dataset, with the human canonical HSPB1 sequence as the reference.

To assess the prevalence of the interaction between the _6_VPFSLL_11_ and _179_ITIPV_183_ motifs and the lateral grooves of human HSPB1, first, the intra and interchain interactions in the 24-mer oligomer of human HSPB1 (PDB ID 6DV5) were mapped using the RING web server [30]. Subsequently, structural models of the human HSPB1 dimer were generated using AlphaFold 2 [22] on a local installation with A30 GPUs. The canonical sequence of human HSPB1 was used as input, along with the UniRef90, MGnify, and BFD sequence databases, and the PDB70 structural database. A total of 500 relaxed structures were generated (5 models with 100 predictions each) in PDB format. The files containing the structures were reformatted using pdb4amber into a PDB format compatible with the ProLIF toolkit [31] and stacked into a single file for processing with CPPTRAJ, which was also used to calculate the RMSD with respect to the reference PDB structure 4MJH, a dimer of the ACD of human HSPB1 in complex with a C-terminal peptide, solved by X-ray diffraction at 2.60 Å [32]. Both tools are part of the AmberTools package [33]. The combined file was then analyzed with the ProLIF toolkit to obtain interaction fingerprint between the residues of the _6_VPFSLL_11_ and _179_ITIPV_183_ motifs and those forming the β4 (_109_LTVKT_113_) and β8 (_153_VSSSL_157_) strands. A contact was defined based on ProLIF’s detection of interactions between any residue in the motifs and any residue in the β4 or β8 strands. Per residue pLDDT values were extracted from the PDB files of the relaxed models using custom Python scripts. Structural models were visualized using VMD [34].

Next, to observe the contacts inferred by AF2 based solely on the multiple sequence alignment (MSA) information, an additional set of 5 models of the HSPB1 dimer was generated, with 5 predictions per model (25 structures), without using templates from the PDB70 database. The distance histograms (distograms) information generated by AF2 for each model along with the Predicted Aligned Error were extracted from the .pkl files using the dgram2dmap tool [35].

## Results

### HSPB1 orthologs dataset characterization

The curated dataset comprises 474 protein sequences, 199 are from invertebrates and 275 from vertebrates. During the curation process and after applying coverage and identity filters, no sequences from other kingdoms, including Plantae, Fungi, Eubacteria and Protista, remained in the dataset. Among the vertebrates, the dataset includes 88 mammals, 124 fish, 26 birds, 26 reptiles, and 11 amphibians. The MSA analysis shows that the ACD is highly conserved in length, with an average of 77.0 ± 0.5 amino acids, while the NTR (88.8 ± 18.3) is longer and more variable compared to the CTR (34.7 ± 8.4). The CTR lengths are similar between vertebrates (32.1 ± 8.1) and invertebrates (38.2 ± 7.5). In contrast, the NTR shows greater length variability between vertebrates (99.5 ± 16.6) and invertebrates (74.0 ± 6.5). This difference is due to a low-complexity region of variable length present in vertebrates, known as the inserted segment in human HSPB1 [13].

The MSA of the full dataset also reveals that the composition of the ACD is highly conserved across all organisms. This pattern is illustrated in Supplementary Figure S1, where the percentage of each type of amino acid in the ACD is similar for most amino acids in both the vertebrate and invertebrate sets. The comparison between the amino acid composition of the NTR and CTR indicates that the NTR contains a higher proportion of aromatic amino acids (TRP, TYR, and PHE), which are nearly absent in the CTR; this enrichment, although unusual for intrinsically disordered regions, is commonly associated with molecular recognition regions that facilitate protein-protein interactions [36,37]. The NTR shows greater variability in amino acid composition between vertebrate and invertebrate sets, with a notable presence of hydrophobic and positively charged amino acids, especially in vertebrates. In contrast, the CTR exhibits a more uniform composition across different organism sets, with a higher proportion of polar and negatively charged amino acids, which may help maintain the protein in solution in the dimeric context. Although both regions are disordered, they exhibit distinct compositions that are linked to their specific roles. It is noteworthy that human HSPB1 contains a single cysteine in the ACD (C137) which, when oxidized, leads to the formation of a disulfide bridge that helps keep the dimer bonded, as shown by experimental data (PDB structure 2N3J) [38]. Given the ability of cysteines to form disulfide bonds that can influence the oligomeric equilibrium of sHSPs, it is intriguing that, despite their low proportion, some organisms have cysteines in the NTR and CTR in addition to the ACD.

### NTR and CTR motifs that interact with the ACD are conserved across all HSPB1 orthologs

We further explored the composition of motifs in the NTR and CTR regions that interact with the lateral grooves of the ACD and play a crucial role in the self-regulation mechanism of HSPB1’s chaperone activity. In human HSPB1, Clouser et al. [13] reported that the _6_VPFSLL_11_ motif in the distal segment of the NTR interacts with the lateral grooves of the ACD formed by β4 and β8 sheets. Similarly, the _179_ITIPV_183_ motif, the extended form of the I/V-X-I/V motif, also interacts with these lateral grooves in the dimeric context [32]. This interaction is also essential for the assembly of oligomers, as demonstrated in the 24-mer structure solved by X-ray crystallography at 3.6 Å and reported in the Protein Data Bank [5]. Compositional data is shown in Supplementary Table S1. The _6_VPFSLL_11_ and _179_ITIPV_183_ motifs show distinct patterns between vertebrates and invertebrates. An essential aspect of this analysis is the use of proline 7 in the NTR, which is fully conserved across both vertebrates and invertebrates, as a reference point for defining the position of the _6_VPFSLL_11_ motif in invertebrates. This conservation guides the alignment and allows the comparison of this motif across species. In vertebrates, the _6_VPFSLL_11_ motif itself occurs in 27.6% of sequences, while the three most frequent alternatives VPFSLL, VPFTFL and IPFTLL together account for 62.9% of sequences.

The _179_ITIPV_183_ motif in the CTR is highly conserved among vertebrates, with alternatives of the I/V-X-I/V motif present in 98.5% of the recruited sequences. In vertebrates, the extended form ITIPV is predominant, found in 56.0% of sequences. Notably, the non-extended form TTIPV occurs in 20.4% of sequences and is exclusively observed in fish species. In contrast, invertebrates display only non-extended forms of this motif. Proline 182 (human HSPB1 numbering) is conserved in 87.5% of sequences, highlighting its critical role in maintaining proper motif conformation during interaction with the ACD. As previously mentioned, mutations at this position (P182A and P182L), are associated with Charcot-Marie-Tooth disease, indicating its importance in HSPB1’s structure and function [15–20].

The analysis of the lateral grooves formed by the β4 and β8 strands within the ACD highlights significant conservation of key amino acids. In vertebrates, in the β4 strand, leucine, valine, and lysine are the predominant amino acids, while in the β8 strand, valine, serine, and leucine are the most frequent ones. Although the composition of the β4 and β8 strands in invertebrates shows greater variability, the residues primarily vary by others with chemically equivalent or similar side chains, indicating that the structural characteristics of the lateral grooves are preserved. This analysis indicates that the composition of the lateral grooves is highly conserved, and despite being located in disordered regions, the motifs interacting with the ACD are also highly conserved or exhibit variants that may fulfill roles similar to those of the _6_VPFSLL_11_ and _179_ITIPV_183_ motifs in human HSPB1. Therefore, the interactions relevant for the self-regulation of human HSPB1 might be characteristic of all HSPB1 orthologs.

### Disordered regions show faster evolution than conserved ACD domain

As it has been mentioned above, the current evolutionary hypothesis for the sHSP family suggests that the disordered regions of small heat shock proteins evolved independently and at a faster rate than the ACD due to the lower structural constraints on these flexible regions [21]. To test this hypothesis in our HSPB1 dataset, we separated the sequences into three distinct regions: NTR, CTR, and ACD, and calculated the evolutionary distances for each region independently. This separation allows us to understand the evolutionary pressures acting on each region of the protein. Since the three datasets were derived from the complete set of orthologs sequences, the evolutionary time that has elapsed between the organisms is equivalent across all three sets. Thus, by measuring the evolutionary distances between distinct regions of the same organism pairs, we can determine whether the evolutionary rates of these regions differ. If the entire protein were subject to the same evolutionary constraints, we would expect a similar distribution of evolutionary distances across all regions. Our analysis of the evolutionary distance distributions revealed distinct patterns for each region. As shown in Figure 2, the probability density functions for the evolutionary distances of the NTR, CTR, and ACD display significant differences, supporting the notion that these regions have evolved at different rates. The distribution for the ACD (green curve) is sharply peaked around low evolutionary distances, indicating that this domain is highly conserved across species. This narrow distribution can be attributed to the structural constraints that limit the variability of the ACD, maintaining its conserved structure.

**Figure 2.**
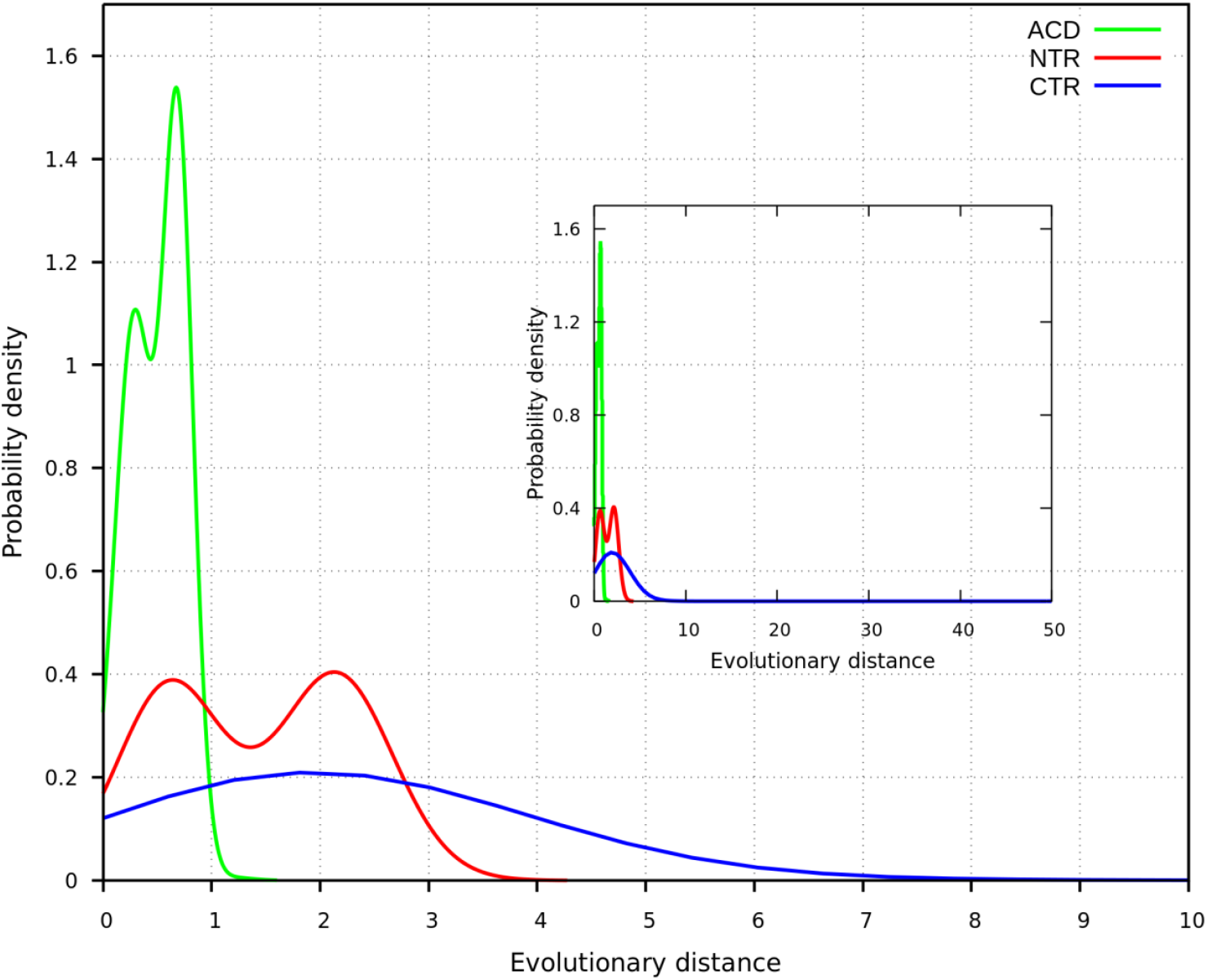
Probability densities of evolutionary distances calculated for the disordered regions and the ACD, with the inset displaying the full range of distance values.

In contrast, the distributions for the NTR and CTR are broader, indicating higher evolutionary variability. The wider spread of these distributions suggests that these disordered regions are less constrained and have accumulated more sequence changes over time, leading to faster evolutionary rates. Additionally, the NTR distribution is slightly shifted towards lower distances compared to the CTR, indicating that, although both regions have evolved faster than the ACD, the NTR may be under slightly stronger evolutionary pressure than the CTR.

Interestingly, the distribution for the NTR appears bimodal. This pattern reflects the distinct evolutionary pressures on the NTR in vertebrates and invertebrates. The first peak, associated with shorter evolutionary distances, corresponds to intra-group comparisons within vertebrates and invertebrates, suggesting recent divergence within each group. The second peak, reflecting higher evolutionary distances, arises from comparisons between vertebrate and invertebrate sequences, indicating greater divergence between these lineages. A similar pattern is observed in the ACD curve, though the distance between the peaks is smaller, highlighting the higher conservation of this domain across species. The inset within Figure 2 expands the view to show the full range of evolutionary distances. While the main peaks of the distributions fall within a limited range, there are some sequences in the CTR with significantly higher distances, indicating more extensive divergence in that region.

The statistical analysis, as shown by the Kolmogorov-Smirnov test, supports these observations. The KS statistic values for the comparisons between NTR and ACD, CTR and ACD, and NTR and CTR are 0.60, 0.80, and 0.31, respectively, with p-values < 1e-16 for all comparisons. These results confirm significant differences in the evolutionary rates of these regions. The broader and more right-shifted probability density distributions for the NTR and CTR, compared to the ACD, indicate that these regions have accumulated more evolutionary changes. Specifically, the higher evolutionary distances observed for the CTR imply that it has experienced the fastest rate of evolution, followed by the NTR, with the ACD being the slowest.

### Motifs in disordered regions interacting with the ACD show reduced evolutionary rates

In light of the previous results, we further explored the evolutionary rates per position of the human HSPB1 sequence using the full dataset of orthologs sequences. The estimated rate profile presented in Figure 3 reveals a clear distinction in the rates of evolution across different regions of the protein. The ACD is characterized by a predominance of positions with low evolutionary rates, which could be attributed to structural constraints that act on this region to preserve its structure, including the β4 (_109_LTVKT_113_) and β8 (_153_VSSSL_157_) strands that frame the lateral grooves (Figure 4.A). However, the low evolutionary rates observed in the β4 and β8 strands likely result from a combination of these structural constraints and functional requirements. According to experimental results, these strands interact with the _6_VPFSLL_11_ motif in the NTR and the _179_ITIPV_183_ motif in the CTR, as well as co-chaperones and client proteins [3,39]. Moreover, this dual role, structural and functional, explains the significant conservation seen in these regions, as supported by the compositional data presented in Supplementary Table S1.

**Figure 3.**
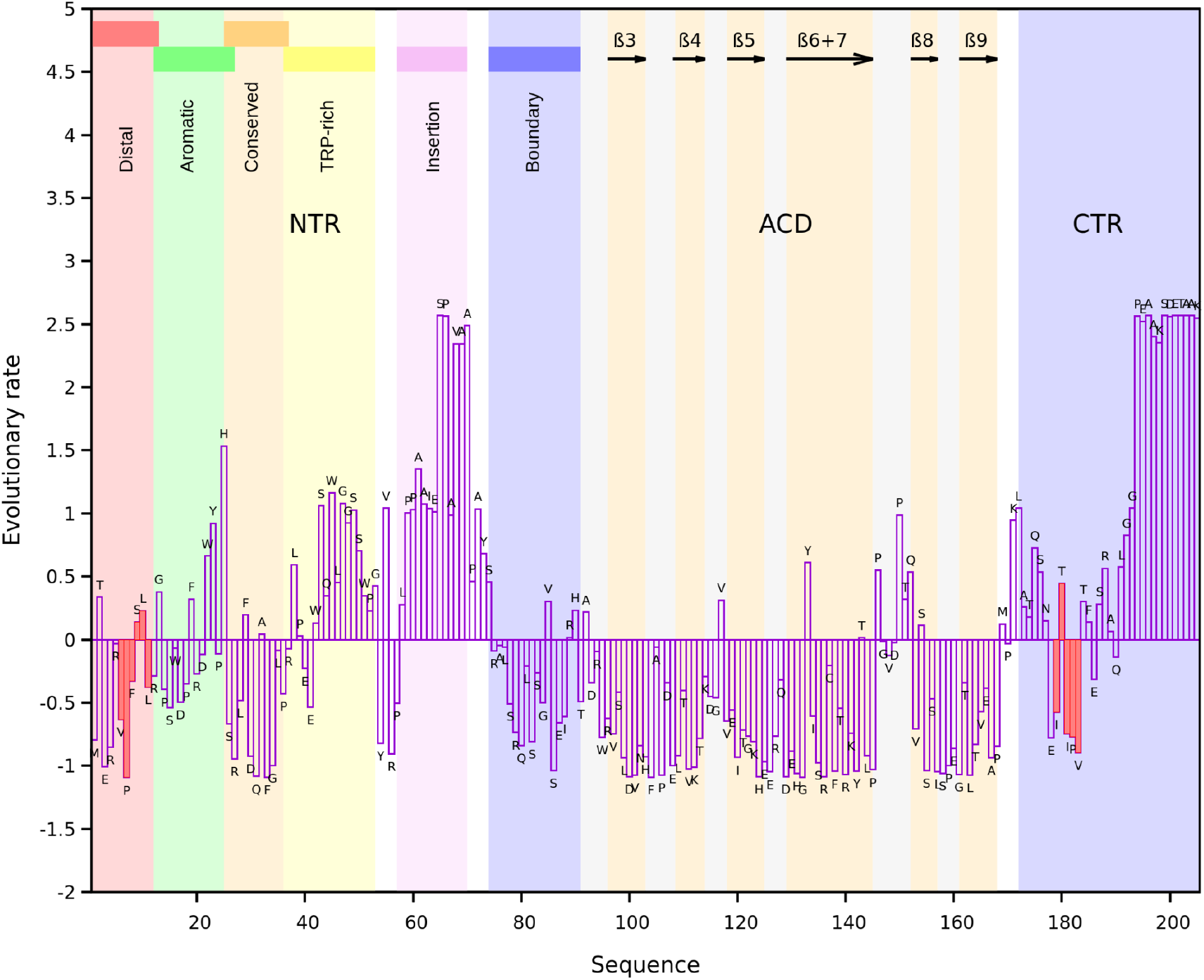
Per position evolutionary rate profile for the canonical human HSPB1 sequence estimated using the full orthologs dataset. The background highlights the NTR, ACD and CTR. The NTR is further divided into segments distal (residues 1-13), aromatic (residues 12-27), conserved (residues 25-37), TRP-rich (residues 37 to 53), insertion (residues 57-70) and boundary (residues 74-91), according to Clouser et al. [13]. The β strands of the ACD are indicated by black arrows. The _6_VPFSLL_11_ motif of the NTR and the _179_ITIPV_183_ motif of the CTR are highlighted in red.

**Figure 4.**
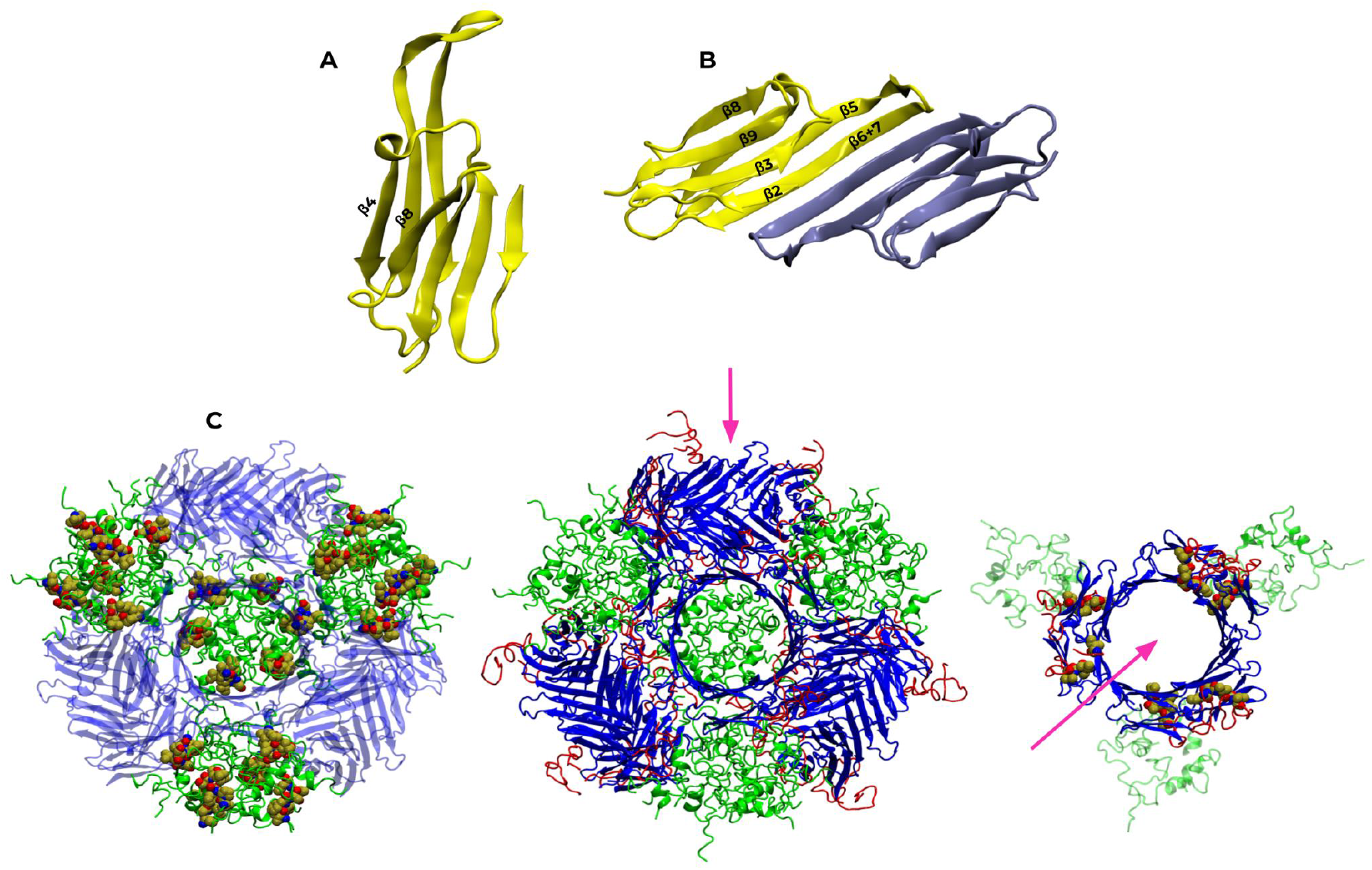
Structural overview of human HSPB1 highlighting the ACD and its oligomeric assembly. **A**. ACD structure of human HSPB1 (PDB ID 4MJH, chain A), showing β4 and β8 strands forming the lateral groove. **B**. Top view of the ACD dimer, indicating the positions of the β-strands. **C**. The central structure shows the 24-subunit oligomeric structure (PDB ID 6DV5) displaying four circular arrangements of three dimeric ACDs (blue), held together through extensive interactions mediated by the NTRs (green). The CTRs of each subunit within the dimers (red) are in contact with the lateral grooves of neighboring subunits. The pink arrow indicates the perspective of the top view on the right, showing one of the four arrangements of six subunits that make up the oligomer. This structure highlights the _179_ITIPV_183_ motif of each subunit interacting with the lateral groove of the adjacent subunit. The left structure emphasizes the arrangement of NTRs in the oligomer, highlighting the _6_VPFSLL_11_ motif, which in this configuration does not interact with the ACD.

The NTR and CTR exhibit a more heterogeneous pattern of evolutionary rates compared to the ACD. Figure 3 shows the rates of the different segments of the NTR (distal, aromatic, conserved, Trp-rich, insertion, and boundary), as defined by Clouser et al [13]. In their study, a phosphomimetic construct was used, incorporating the mutations S15D, S78D, and S82D in the NTR to mimic the phosphorylation state of human HSPB1, and substituting the _179_ITIPV_183_ motif in the CTR with _179_GTGPG_183_. This construct allowed the study of the dimeric form of human HSPB1 due to its inability to form oligomers larger than dimers. Hereafter, any reference to experimental results related to the dimeric form will pertain to this construct.

Regarding the interactions between the NTR and the ACD in the dimer, the segments of the NTR can be divided into two groups: those that interact with the ACD and those that do not. Among the interacting segments, the distal segment contains the _6_VPFSLL_11_ motif, which engages with the lateral grooves of the ACD. The conserved segment includes the _26_SRLFDQXFG_34_ motif, which interacts with the central groove formed at the dimer interface. The aromatic segment in human HSPB1 has also been shown to interact with the ACD, and in vertebrates, this region contains alternative sequences of a characteristic motif present in the NTR of sHSPs, _16_WDPF_19 [40],_ involved in chaperone oligomerization [41,42]. Although the canonical form of this motif is not conserved in the recruited invertebrate sequences, the corresponding region in the MSA is enriched with tryptophan and negatively charged residues. Notably, 68.3% of these sequences exhibit the specific sequence D/N-W-W-D/E, highlighting the importance of the chemical properties of these amino acids in this segment of the NTR. Additionally, the boundary segment has been demonstrated to form an antiparallel β-sheet (β2) with the β3 strand of the ACD within the same chain (Figure 4.B) [13]. These four segments show lower than the average evolutionary rates, likely due to the functional importance of their interactions with the ACD, despite being located in a disordered region. In contrast, the Trp-rich and insertion segments do not interact with the ACD in a sustained manner in the dimeric form and exhibit evolutionary rates higher than the average of the full sequence. In particular, the insertion segment behaves as a solvated random coil [13], with its length and composition varying significantly among different organisms. This variability may account for the higher evolutionary rates than average observed in this region, as solvent-exposed regions tend to evolve more rapidly compared to those with lower accessible surface areas [43].

The CTR generally exhibits evolutionary rates above the sequence average, except in positions adjacent to and within the extended I/V-X-I/V motif, _179_ITIPV_183_, where rates fall below this average. This pattern of conservation within the motif and its surrounding residues underscores the functional importance of the motif’s context. Structural analyses indicate that adjacent residues enhance binding specificity, complementing the motif’s core binding role [44]. Additionally, the crystal structure of a dimeric construct containing only the ACD of human HSPB1 (PDB ID 4MJH), complexed with the _179_ITIPVTFE_186_ peptide from the CTR, shows this peptide, which includes the motif of interest, interacting with the lateral grooves of the ACD [32]. Evolutionary rates of residues within the _6_VPFSLL_11_ and _179_ITIPV_183_ motifs display distinct behaviors (positive or negative rates), suggesting differential tolerance to substitutions. Within the _179_ITIPV_183_ motif in human HSPB1, the variable positions T180 and P182 exhibit opposite evolutionary rates: T180 has a positive rate, whereas P182 shows a negative rate. Notably, invertebrates and fish lack the extended form seen in human HSPB1, which includes T180, preserving only the final three positions of the motif across all analyzed species.

To analyze the relationship between the evolutionary rate values for different positions or regions of human HSPB1, it is necessary to consider not only the contacts involving various regions at the dimeric level but also the contacts implicated in the formation of larger oligomers, as detailed in Supplementary Tables S2-S6. Intra and interchain contacts within the dimer, as well as the interchain contacts implicated in the higher-order oligomer stabilization, play key roles in regulating the chaperone activity of human HSPB1. The 24-subunit oligomeric structure of human HSPB1 reported in the Protein Data Bank is arranged into four groups of three dimer pairs connected through extensive NTR interactions (Figure 4.C). Residues corresponding to the inserted segment and part of the boundary segment were not solved in this PDB structure due to their high mobility [5]. Analysis of the interactions that take place in this structure shows that, unlike in the dimer where the _6_VPFSLL_11_ motif interacts with the ACD’s lateral grooves, in the oligomeric structure it is incorporated into the NTR arrangements that hold the four groups together. These arrangements account for the 57.5% of the interchain interactions within the structure (Supplementary Table S2). A total of 90 interchain contacts were mapped for the _6_VPFSLL_11_ motif in the 24 subunits, which are not symmetrical. Predominantly the interactions are of Van der Waals type (Supplementary Table S3). F8 and L11 represent the greatest part of the interactions (47 and 21 respectively), and 14 chains are engaged through a π-π stack between F8 and tryptophans that belong to the TRP-rich segment (Supplementary Table S5).

While the NTRs do not account for interchain interactions with the ACD in the 24-mer structure, the contacts between the CTR and the ACD represent 10.5% out of the total of interchain contacts mapped (Supplementary Table S2). In this structure, each subunit engages the lateral groove of the neighboring one, positioning the _179_ITIPV_183_ motif within this groove. A total of 29 interactions between this motif and the β4 and β8 strands were mapped, including 21 of Van der Waals type and 8 H-bonds (Supplementary Table S4). Extending the analysis to the neighboring residues encompassed in the _179_ITIPVTFE_186_ peptide crystallized in the 4MJH structure, increases the number of interactions up to 50, mainly due to Van der Waals or π-H-bond interactions between F185 and S155 from β8, that is found to occur in all chains (Supplementary Table S6), highlighting the relevance of these neighboring residues for surface binding. Nevertheless, it must be taken into account that, although the PDB structure constitutes a valuable source of information about the complex topology, it only captures a single conformation of the oligomer. Thus, the number and nature of the interactions mapped in this structure might differ in a dynamical context. In this line, the work of Clouser et al. reports that in human HSPB1 oligomers the CTRs containing the _179_ITIPV_183_ motif can exist both in a free state and bound to the grooves formed by β4/β8 strands [13].

The significance of the interactions between the _6_VPFSLL_11_ and the _179_ITIPV_183_ motifs and the lateral grooves of the ACD can also be understood considering additional information provided by experiments. It has been observed that phosphomimetic mutations alone (S15D, S78D, and S82D) do not prevent oligomerization, as the _179_ITIPV_183_ motif in the CTR still interacts with the ACD’s lateral grooves [8]. Consequently, substituting _179_ITIPV_183_ with _179_GTGPG_183_ is essential to isolate a dimeric form, establishing this construct as a standard reference for studying the chaperone activity of human HSPB1 by reliably preventing higher-order oligomer assembly. Furthermore, structural data from all oligomeric sHSPs reported in the PDB consistently show that the I/V-X-I/V motif, conserved in most sHSPs, forms intersubunit interactions with the lateral grooves of the ACD regardless of the oligomer topology [10,45,46]. This prevalence likely reflects the critical functional and structural role of this interaction, as evidenced by the lower than average evolutionary rates observed for residues comprising the motif, indicative of the constraints acting to maintain this interaction. Experimental evidence also shows that a human HSPB1 construct lacking the distal segment of the NTR (residues 1 to 14), which includes the _6_VPFSLL_11_ motif, can still form oligomers, although they are smaller than those of the wild type form [47].

In contrast to the prevailing interactions between the CTR and the lateral grooves of the ACD observed in PDB structures of sHSP’s of different size, the arrangement of NTRs differs significantly among these, depending on how the ACDs of each chain interact to form the dimers that constitute the building blocks of higher-order oligomers [1,46]. Observed arrangements include those where the NTRs may face inward toward the core, outward, or sideways [10]. Additionally, oligomer polydispersity is a major factor that must be considered from this structural standpoint. For instance, a study on HSPB1 from Chinese Hamster, sequence included in the orthologs dataset, shows that its quaternary structure varies according to temperature and concentration, forming oligomers of different sizes that coexist in a dynamic equilibrium, where specific conditions favor the formation of smaller or higher order oligomers [48].

Thus, considering the polydispersity of oligomers and the diversity of NTR arrangements across known oligomeric structures of sHSPs from different organisms, as well as the variability in NTR length among sequences in our HSPB1 orthologs dataset (primarily due to the variable length of the inserted segment in vertebrates and its absence in invertebrates), and the lack of PDB structures for these proteins, it would be uncautious to interpret the role of the _6_VPFSLL_11_ motif in oligomerization and its contribution to the evolutionary rates of its residues due to this process solely based on the available oligomeric structure of human HSPB1. Nevertheless, the fact that oligomer formation in human HSPB1 still occurs in the absence of the distal segment comprising this motif indicates, as observed in the mapped contacts within the 24-mer structure, that oligomerization does not rely exclusively on this region [47]. The experimental evidence showing that the interaction between the _6_VPFSLL_11_ motif and the lateral grooves of the ACD regulates chaperone activity in the human HSPB1 dimer [14], along with the prevalence of this motif or its variants in the sequences of the recruited orthologs, support the idea that these positions likely exhibit low evolutionary rates due to selective pressures to preserve the interaction between the motif and the lateral grooves of the ACD at the dimeric level.

### Structural modeling of human HSPB1 with AlphaFold 2 reveals insights on coevolution

We generated 500 structural models of the human HSPB1 dimer using AF2. As mentioned above, this approach was used to evaluate the occurrence of interactions between the _6_VPFSLL_11_ and _179_ITIPV_183_ motifs, located in the disordered regions, and the β4 and β8 strands that frame the lateral grooves. Contacts involving the _179_ITIPV_183_ motif and the lateral grooves have been observed in the crystallographic structure 6DV5, which reveals the interaction between this motif and the lateral grooves of the ACD within the oligomeric structure. Although structural data is available for the NTR in the phosphomimetic form described by Clouser et al. [13], there is currently no available structure exhibiting the interaction between the _6_VPFSLL_11_ motif and the lateral grooves in the canonical form of HSPB1. Therefore, we used AF2 to investigate whether this interaction could also be predicted in the canonical structure.

Structural analysis indicated that the structure of the ACD (residues 92 to 168) was conserved across all models (RMSD for backbone atoms was 0.51 ± 0.15, using the 4MJH PDB structure as a reference), while the disordered regions adopted various conformations (Supplementary Figure S2). The average per residue estimate of confidence (pLDDT) exhibited values above 70 with low standard deviation for the residues forming the ACD, which indicates that this region is well modeled (Supplementary Figure S3). In particular, positions with values close to or exceeding 90 suggest high accuracy in the models. In contrast, pLDDT scores for the residues forming the CTR were below 50, as expected due to their disordered nature [49]. On the other hand, the high pLDDT scores for the distal segment of the NTR containing the _6_VPFSLL_11_ motif, align with AF2’s capacity to identify conditionally folded intrinsically disordered regions. This interaction-driven structural stabilization of the NTR is likely reflected by the coevolutionary signals captured in the multiple sequence alignment used by AF2 [50].

From the interaction fingerprint obtained from the model set, we assessed whether any of the motifs mentioned were in contact with the lateral grooves of the ACD. The analysis was performed separately for each chain, and the contacts present were found to be symmetric between the two chains. This revealed that in 98.2% of the predicted structures, the distal segment of the NTR containing the _6_VPFSLL_11_ motif, was in contact with one of the lateral grooves of the dimer through Van der Waals and H-bond interactions (Figure 5). In the remaining 1.8%, the NTR was not in contact with the lateral grooves but instead adopted an extended conformation. None of the models showed the _179_ITIPV_183_ motif of the CTR interacting with the grooves. For the _6_VPFSLL_11_ motif, 68.6% of the interactions were intrachain contacts, while 29.6% were interchain contacts. Both types of interactions are consistent with the “quasi-ordered” states that human HSPB1 can adopt [13].

**Figure 5.**
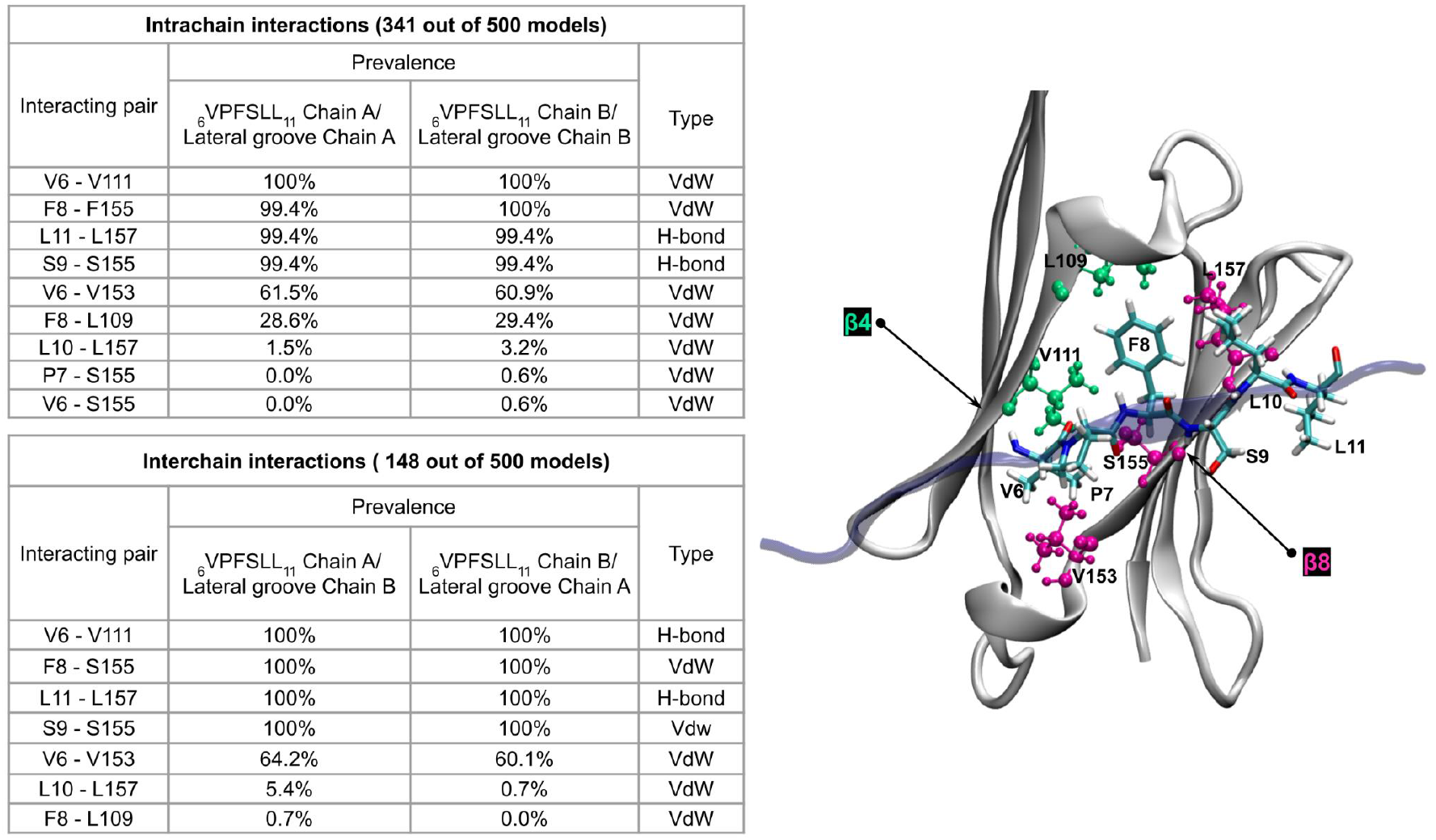
Summary of intrachain and interchain interactions between the _6_VPFSLL_11_ motif and the lateral grooves of the ACD (β4 and β8 strands) in human HSPB1 dimer models. The prevalence of interactions is shown as a percentage of the total number of models in which each type of contact (intra or interchain) occurs, along with the type of contact (Van der Waals or H-bond). The right panel highlights the residues involved in the interactions listed in the table.

Analysis of the interacting residue pairs in human HSPB1 dimer models reveals that the F8-S155 pair interacts consistently in nearly 100% of models displaying contacts, regardless of whether these interactions are intra or interchain. This observation aligns with findings by Bauggman et al., who reported that substitutions at these specific positions (F8G within the _6_VPFSLL_11_ motif and S155Q in the β8 strand) in a phosphomimetic construct of human HSPB1 enhanced its chaperone activity toward the natural client tau [14]. The authors attributed this increase in activity to the release of the NTR from the β4-β8 lateral groove, suggesting that this displacement allows the NTR to adopt conformational states better suited for effective tau binding. The broader relevance of this mechanism is further supported by a previous study on HSPB5 [12], where a comparable groove mutation (S135Q) similarly enhances chaperone activity toward amyloid-β, implying that NTR release may serve as a general activation mechanism across sHSP-client interactions.

In a complementary analysis, we generated 25 additional models using AF2 configured to rely exclusively on MSA data. This approach provided an opportunity to observe how AF2 infers contacts relying only on sequence-derived evolutionary signals. Throughout the recycling process, AF2 generates distograms as intermediate outputs, which predict the probabilistic distances between residue pairs and help refine the model’s structural predictions. Although these distograms are not final outputs, they provide insight into the algorithm’s assessment of inter-residue relationships. In these template-free models, the dimer’s ACD structure was conserved, except in one model where the quaternary structure of the ACD did not align with the 4MJH PDB. Although each chain’s individual structure was preserved, the quaternary arrangement diverged significantly, as evidenced by high Predicted Aligned Error values, which specifically indicated a lack of confidence in the interchain contacts and the overall quaternary structure alignment. As a result and the low occurrence of this problem (1/25 models), this model was excluded from subsequent analyses.

The distogram shown in Figure 6 is an averaged representation of these models, indicating predicted distances between residue pairs in the human HSPB1 dimer. Residues 1-91, 92-168, and 169-205 of Chain A, and their respective counterparts in Chain B (206-410), represent the NTR, ACD, and CTR, respectively. The ACD forms the central block, with shorter intrachain distances (∼5 Å) shown in green, reflecting conserved interactions within each ACD. The purple rectangles highlight the β6/7 strands (133-142), forming interchain contacts, also around ∼5 Å, consistent with the structured and conserved nature of the ACD dimer interface. Residues belonging to the CTR do not appear to establish contacts with the ACD. In contrast, the distal segment of the NTR of each chain (residues 1 to 13), which contains the _6_VPFSLL_11_ motif, and the β4 and β8 strands (residues 109 to 113 and 153 to 157, respectively) that frame the lateral grooves, are predicted to be in contact. Intrachain contacts between this segment and each strand are marked by black rectangles, with distances ranging from ∼10 Å to 15 Å. Interchain contacts, highlighted in blue, show distances of ∼15 Å to 20 Å. These values suggest transient or flexible associations rather than direct atomic interactions.

**Figure 6.**
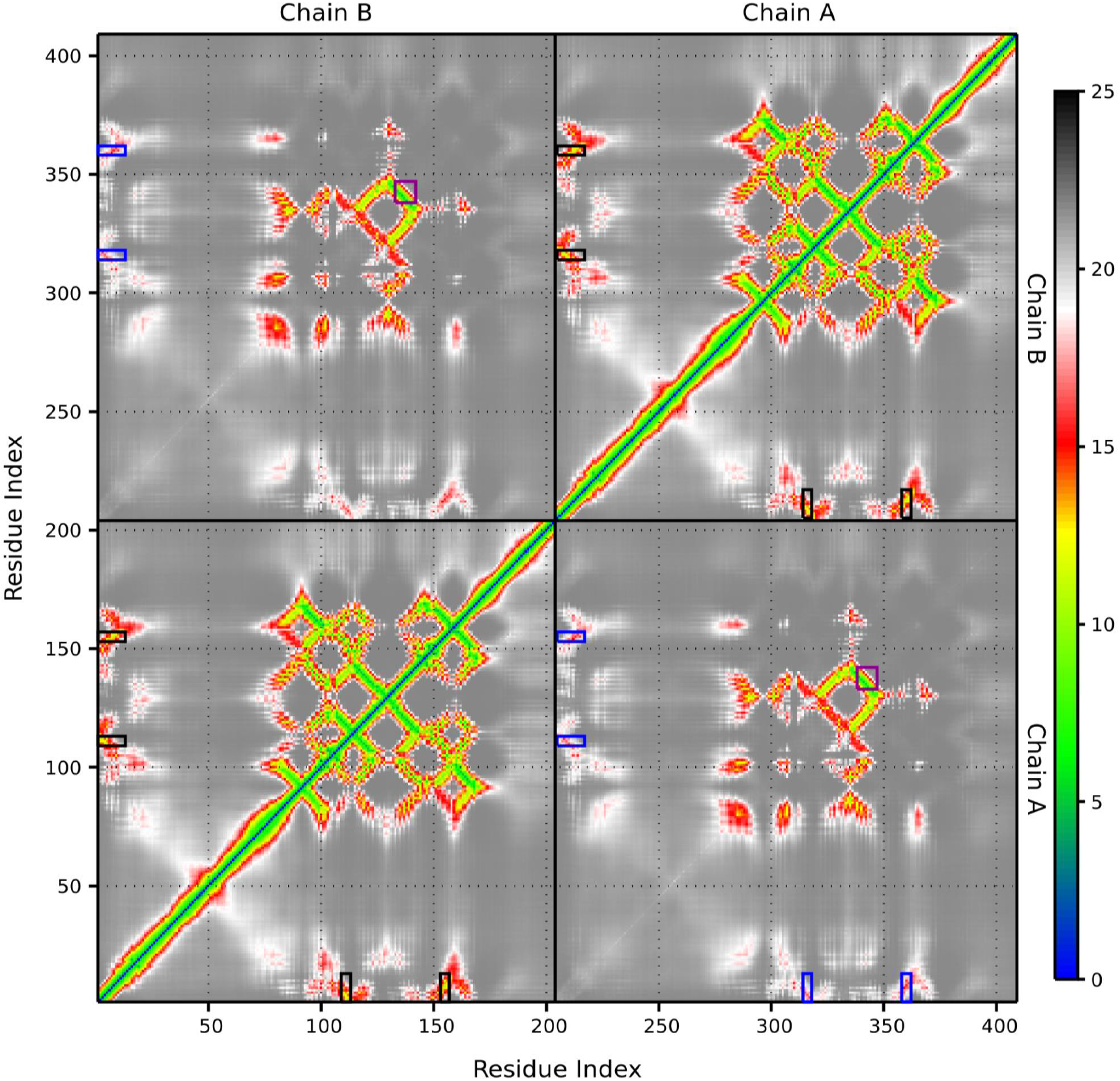
Average distogram generated from individual distograms produced by AF2 for human HSPB1 dimer models without the use of the PDB70 database. Colors represent the probability of contact between residue pairs, with blue indicating shorter distances and red indicating longer distances. Black and blue rectangles highlight the intrachain and interchain contacts, respectively, between the distal segment containing the _6_VPFSLL_11_ motif and the β4 and β8 strands that form the lateral grooves. Purple rectangles emphasize the intersection of the β6/7 strands (residues 133-142) from each chain, forming the dimeric interface. Dividing lines denote the areas corresponding to each chain, with residue index ranging from 1 to 205 for Chain A and 206 to 410 for Chain B.

It is important to note that these distances are not final, as seen in the relaxed models used for evaluating contact occurrence with Prolif. Instead, they are intermediate predictions during the recycling process, reflecting AF2’s ability to capture flexible or transient associations based on coevolutionary signals from the MSA, even without template guidance. Figure 6 was generated by averaging the distograms from individual models, with each point including the standard deviation. The standard deviation for the ACD region remains close to 0, indicating consistent predictions, whereas the NTR-lateral groove contacts exhibit deviations up to 2.5 Å, reflecting greater flexibility in these regions (data not shown).

## Discussion

In this work, we explore the implications of the interactions between _6_VPFSLL_11_ motif of the NTR and _179_ITIPV_183_ of the CTR with the ACD in the evolution of human HSPB1. These contacts play a key role in the self-regulation of chaperone activity: the _6_VPFSLL_11_ motif has a regulatory role at dimeric level, while the _179_ITIPV_183_ motif is implicated in oligomer formation [13,14,51]. The current evolutionary hypothesis for the sHSPs superfamily suggests that the NTR and CTR evolved independently and at different rates than the conserved ACD [21]. Since residues involved in functionally relevant interactions are often evolutionarily constrained and tend to coevolve to maintain their roles [52], we conducted evolutionary analysis and structural modeling to assess coevolution between the disordered regions and the ACD of human HSPB1. The results were analyzed in light of structural and functional information reported in the literature, and the occurrence of disease-related variants at positions implicated in these contacts was also considered.

To understand the impact of these interactions on the evolution of human HSPB1 and its role in chaperone activity, we worked with a manually curated set of orthologs sequences of human HSPB1. This dataset revealed that the amino acid composition of the ACD is conserved in both vertebrates and invertebrates, while the NTR and CTR exhibit greater variability. MSA analyses indicate that the composition of the lateral grooves of the ACD is highly conserved, and that the positions of the _6_VPFSLL_11_ and _179_ITIPV_183_ motifs are either preserved in orthologs or replaced by amino acids with comparable physicochemical properties, likely maintaining their functional interaction with the ACD. This is consistent with the fact that many motif-binding domains exhibit weak specificity, interacting primarily with a small core of residues while tolerating substitutions that retain essential binding characteristics, thereby allowing critical interactions to persist despite evolutionary divergence [53,54]. To determine if our dataset mirrors the evolutionary pattern reported by Kriehuber et al. [21], despite containing less divergent sequences, we replicated their methodology. Assuming uniform evolutionary constraints would produce similar rates across all regions, our findings instead show distinct rates: the ACD evolves the slowest, followed by the NTR, with the CTR evolving the fastest. Although disordered regions collectively evolve faster than the ACD, individual positions within them do not evolve uniformly. In particular, sites involved in ACD interactions, such as the _6_VPFSLL_11_ and _179_ITIPV_183_ motifs, evolve more slowly than the average of the full sequence, likely due to selective pressures preserving these critical interactions. This also explains why the NTR, despite being longer than the CTR, has a lower overall evolutionary rate, as it contains a greater number of ACD-interacting motifs.

AlphaFold 2 infers coevolutionary-like relationships by leveraging phylogenetic information in MSAs without directly calculating traditional coevolution metrics or explicit covariance statistics [22]. Through this approach, the model implicitly learns inter-sequence relationships that capture evolutionary constraints essential for protein structure, enabling it to predict contacts that mirror coevolutionary events. The generation of structural models for the human HSPB1 dimer captures these critical contacts, and notably, these interactions are also predicted in models generated without the use of templates. This consistency indicates that the coevolutionary-like relationships inferred by AF2 are grounded in evolutionary constraints, supporting their use as evidence of coevolution in our analyses.

Together, this evidence suggests that, although the disordered regions of HSPB1 exhibit higher substitution rates on average compared to the ACD, they have not evolved independently; instead, they have likely coevolved with the ACD to preserve essential interactions necessary for chaperone activity regulation. These findings are consistent with studies on proteins with disordered regions, where evolutionary conservation is often observed in positions or motifs within these regions that participate in contacts with ordered domains [1,55,56]. Moreover, this evolutionary perspective provides a context for understanding why substitutions in conserved motifs within disordered regions, such as P7S and P7R variants in the _6_VPFSLL_11_ motif or the P182A and P182L variants in the _179_ITIPV_183_ motif, are associated with Charcot-Marie-Tooth disease [15–20]. These motifs interact with the ACD’s lateral grooves, and it is likely that these substitutions alter their interaction, potentially affecting human HSPB1’s chaperone function, as even a single amino acid change at a critical site can be enough to disrupt binding [54]. For the P182 residue, it has been shown that although proline does not directly interact with the ACD, it restricts the conformations of the _181_IPV_183_ motif available for interaction with the ACD. Replacing proline with leucine leads to a lower binding affinity between the lateral grooves and the motif, which is attributed to an increase in the conformational flexibility of the motif due to the loss of torsional constraints imposed by proline [15].

Interactions between specific motifs from the disordered regions (I/V-X-I/V variants in the NTR and CTR) have also been observed in other human paralogs [12,32,57], and competition for the lateral grooves between these motifs has been reported for human HSPB5 [58], similar to the interactions seen with _6_VPFSLL_11_ and _179_ITIPV_183_ as described by Clouser et al [13]. Given that various human paralogs feature interaction motifs in the NTR, CTR, or both [3], it is plausible that coevolutionary processes driven by selective pressures to maintain critical interactions could also occur in other members of the sHSP family. Further studies combining structural and evolutionary information as well as the impact on chaperone activity of disease associated variants, could improve our understanding of activity diversity of these sHSPs.

## Acknowledgments

This work used computational resources from CCAD – Universidad Nacional de Córdoba (https://ccad.unc.edu.ar/), which are part of SNCAD – MinCyT, República Argentina. MSF and LRS are researchers and VER is postdoctoral fellow at Consejo Nacional de Investigaciones Científicas y Técnicas (CONICET). FNEB is a PhD student at Universidad Nacional de Quilmes (UNQ). The research was supported by Universidad Nacional de Quilmes (PUNQ 827-2282/22). Agencia Nacional de Promoción Científica y Tecnológica (PICT-2020-0269) and Fundación Florencio Fiorini (Grants for Research in Biomedical Sciences 2024: Effect of pathogenic variants on the self-regulation mechanism of human HSPB1 chaperone activity).

## Author contributions

VER and FNEB obtained, curated, and characterized the HSPB1 orthologs dataset. VER and MSF conceptualized, designed the study, conducted structural and evolutionary analysis and prepared the initial draft. LRS contributed to conceptualization, manuscript revision, and visualization. MSF and LRS provided funding for this work. All authors contributed to discussing the results.

## Competing Interests

The authors declare no competing interests.

## Data, Materials, and Software Availability

The multiple sequence alignment of the complete HSPB1 orthologs sequence dataset is available at the CONICET Digital Institutional Repository - Research Data (http://hdl.handle.net/11336/248901).

## Notes

http://hdl.handle.net/11336/24890

### Competing Interest Statement

The authors have declared no competing interest.

