## Supplementary Figures and Tables for "Coevolution in Small Heat Shock Protein 1 (HSPB1) is Promoted by Interactions between the Alpha-Crystallin Domain and the Disordered Regions"

### Supplementary information

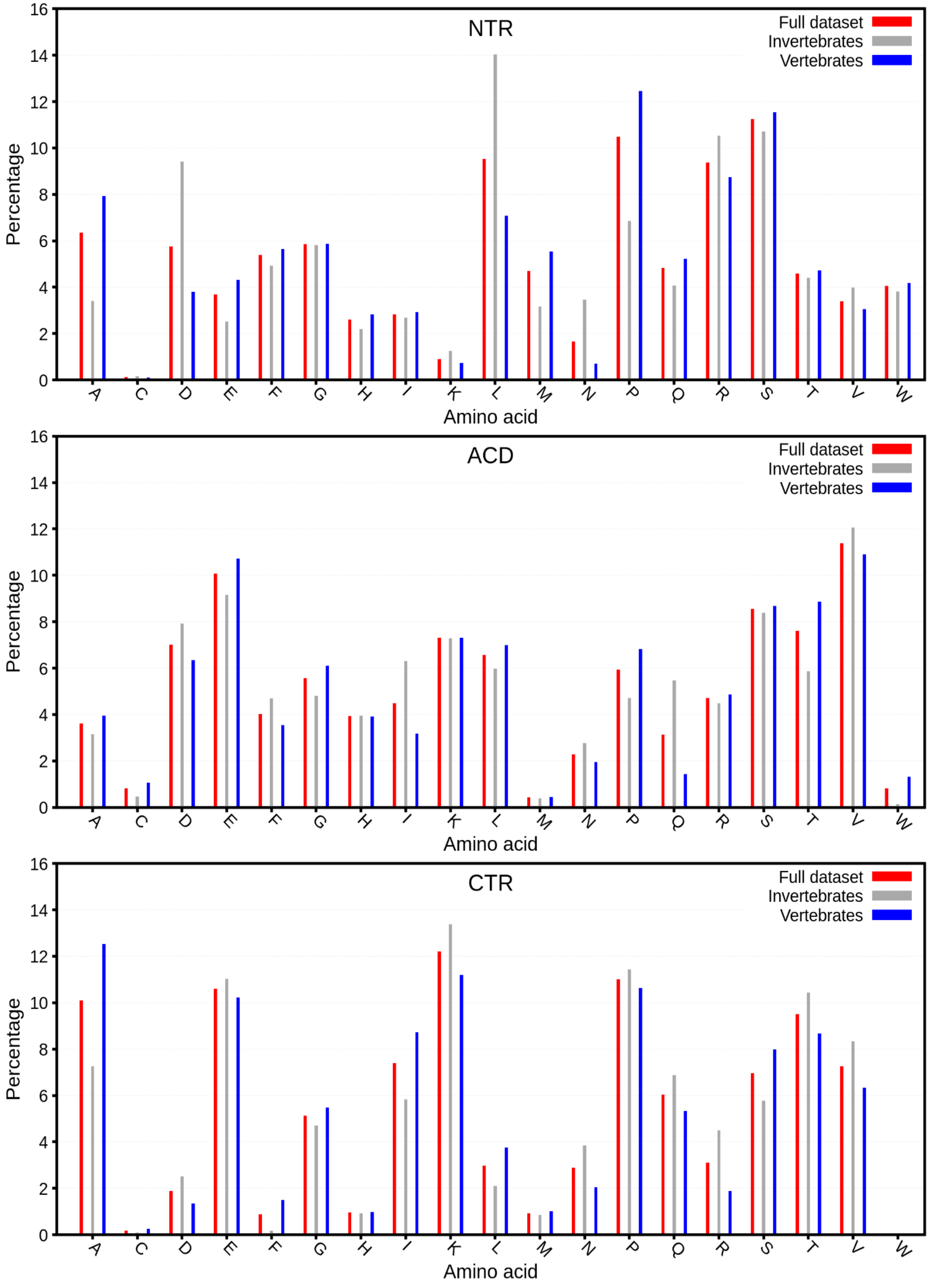


**Figure S1.** Amino acid composition of the NTR, CTR, and ACD in the HSPB1 sequences from the full dataset, as well as the vertebrate and invertebrate subsets.

| Amino acid composition per position of the interacting motifs | | | | |
|---|---|---|---|---|
| Dataset | $_{6}$VPFSLL$_{11}$ | | $_{179}$ITIPV$_{183}$ | |
| | % Composition | Most frequent alternative sequences | % Composition | Most frequent alternative sequences |
| Vertebrates | V V=55.6, I=44.4<br>P P=100.0<br>F F=98.6, L=0.7, T=0.4<br>T T=57.8, S=40.7, A=1.1<br>L L=53.8, F=37.8, M=5.1<br>L L=85.1, M=5.5, Q=4.0 | VPFSLL: 27.6%<br>VPFTFL: 21.5%<br>IPFTLL: 13.8% | I I=66.6, T=24.4 V=6.2<br>T T=82.9, S=5.8, N=5.5<br>I I=98.9, F=0.4, Q=0.4<br>P P=99.3, R=0.3, E=0.4<br>V V=96.4, I=2.6, S=0.7 | ITIPV: 56.0%<br>TTIPV: 20.4%<br>INIPV: 4.4% |
| Invertebrates | V V=66.8, L=22.6, I=5.5<br>P P=100.0<br>L L=61.3 F=15.1, M=11.6,<br>L L=38.2 M=28.6, V=13.1,<br>F F=73.9, L=13.1, Y=6.0<br>- -=98.5, P=1.0, G=0.5 | VPLMF: 19.6%<br>VPLLF: 14.1%<br>VPLVF: 6.0% | V V=72.4, I=27.1, -=0.5<br>P P=70.9, Q=9.6, S=4.5<br>I I=88.4, V=10.6, L=0.5 | VPI: 53.8%<br>IPI: 12.1%<br>VQI: 8.5% |
| Full dataset | V V=60.3, I=28.1, L=9.5<br>P P=100.0<br>F F=63.5, L=26.2, M=4.9<br>T T=34.2, S=23.8, L=16.0<br>F F=53.0, L=36.7, M=3.2<br>L L=49.4, R=25.1, D=4.2 | VPFSLL: 16.0%<br>VPFTFL: 12.5%<br>VPLMFR: 8.2% | I I=68.8, V=30.4, L=0.2<br>P P=87.6 Q=4.0, N=1.9<br>V V=60.3, I=38.6, L=0.2 | IPV: 56.3%<br>VPI: 22.3%<br>IPI: 6.5% |

| Amino acid composition of the lateral grooves | | |
|---|---|---|
| Dataset | $_{109}$LTVKT$_{113}$ (β4 strand) | $_{153}$VSSSL$_{157}$ (β8 strand) |
| | % Composition | % Composition |
| Vertebrates | L L=98.6, I=1.5<br>V V=52.7, T=38.6, M=3.6<br>V V=98.6, I=1.5<br>K K=98.9, R=1.1<br>T T=97.8, M=2.2 | V V=91.3, I=8.4<br>S S=34.9, T=32.7, R=17.1<br>S S=98.6, P=0.7, A=0.4<br>S S=89.8, T=5.1, A=2.9<br>L L=99.6, A=0.4 |
| Invertebrates | I I=94.0, L=4.0, V=2.0<br>T T=60.3, S=21.6 N=7.5<br>V V=99.0, I=1.0<br>K K=97.5, R=2.5<br>T T=62.3, V=25.1, I=7.0 | V V=77.9, I=18.1 L=4.0<br>T T=37.7, V=25.1 E=16.6<br>S S=98.5, C=1.5<br>S S=50.8, T=19.6, R=12.6<br>L L=99.5, I=0.5 |
| Full dataset | L L=58.9, I=40.3, V=0.8<br>T T=47.7, V=31.2, S=9.7<br>V V=98.7, I=1.3<br>K K=98.3 R=1.7<br>T T=82.9, V=10.6, I=3.0 | V V=85.7, I=12.5, L=1.7<br>T T=34.8, S=23.2, V=12.9<br>S S=98.5, C=0.8, P=0.4<br>S S=73.4, T=11.2, R=5.3<br>L L=99.6, I=0.2, A=0.2 |

**Table S1.** Per-position percentage composition of amino acids in the sequence alignment of vertebrates, invertebrates, and the full dataset for the VPFSLL, I/V-X-I/V motifs, and the β4 and β8 strands. For the motifs located in disordered regions, the three most frequent alternative motifs in each alignment are indicated. In each position, only the percentage of the predominant amino acids (up to three) is shown.

| Intrachain interactions (total number of mapped interactions = 2811) | | | |
|---|---|---|---|
| | NTR | ACD | CTR |
| NTR | 34.2% | 4.4% | 2.5% |
| ACD | 4.4% | 48.4% | 3.7% |
| CTR | 2.5% | 3.7% | 6.8% |
| **Interchain interactions (total number of mapped interactions = 830)** | | | |
| | NTR | ACD | CTR |
| NTR | 57.5% | 0.0% | 9.1% |
| ACD | 0.0% | 22.9% | 10.5% |
| CTR | 9.1% | 10.5% | 0.0% |

**Table S2.** Percentage of intra and interchain interactions involving the NTR (residues 1–91), ACD (residues 92–168), and CTR (residues 169–205) in the 24-mer structure of human HSPB1 (PDB ID 6DV5). Values represent the proportion of interactions between and within each region relative to the total mapped interactions.

| Interchain interactions between $_{6}$VPFSLL$_{11}$ motif and NTR residues | | |
|---|---|---|
| **Residue** | **Number of interactions** | **Interaction type** |
| **V** | 4 | Van der Waals |
| **P** | 7 | Van der Waals |
| **F** | 47 | Van der Waals (25) / π-π Stack (11) / π- H-bond (1) |
| **S** | 6 | Van der Waals |
| **L** | 5 | Van der Waals |
| **L** | 21 | Van der Waals |

**Table S3.** Summary of interchain interactions between residues within the $_{6}$VPFSLL$_{11}$ motif and NTR residues from other chains within the 24-mer structure of human HSPB1.

| Interchain interactions | | | | |
|---|---|---|---|---|
| **Residue** | **β4 ($_{109}$LTVKT$_{113}$)** | | **β8 ($_{153}$VSSSL$_{157}$)** | |
| | **Number** | **Interaction type** | **Number** | **Interaction type** |
| **I** | 0 | - | 0 | - |
| **T** | 8 | H-bond (6) / Van der Waals (2) | 0 | - |
| **I** | 0 | - | 0 | - |
| **P** | 3 | Van der Waals | 6 | Van der Waals |
| **V** | 0 | - | 9 | Van der Waals |
| **T** | 0 | - | 3 | H-bond (2)/ Van der Waals (1) |
| F | 0 | - | 20 | Van der Waals (14) / π- H-bond (5)/ H-bond (1) |
| E | 0 | - | 1 | H-bond |

**Table S4.** Interchain interactions in the 24-mer structure of human HSPB1 between the $_{179}$ITIPVTFE$_{186}$ peptide of the CTR and residues in the β4 ($_{109}$LTVKT$_{113}$) and β8 ($_{153}$VSSSL$_{157}$) strands, which frame the lateral grooves of the ACD.

| Residues involved in interchain interactions between $_6$VPFSLL$_{11}$ motif and NTR residues | | | | | | | |
|---|---|---|---|---|---|---|---|
| **Motif** | **AA** | **Chains in contact** | **Type** | **Motif** | **AA** | **Chains in contact** | **Type** |
| V6 | A32 | F R<br>E S | VdW | F8 | W16 | S<br>N | π - H-bond |
| | E40 | K N<br>J O | VdW | | S49 | I<br>P | VdW |
| P7 | F33 | A E O<br>B F N | VdW | | H25 | K<br>R | π - π Stack |
| | Y54 | A M Q<br>B L P | VdW | | P24 | O<br>J | VdW |
| | W42 | W<br>V | VdW | | W51 | C<br>D | π - π Stack |
| F8 | W42 | E C M Q G I W<br>F D L P H X V | π - π Stack | S9 | S15 | B D N<br>Q G S | VdW |
| | | E I<br>F X | VdW | | W16 | X<br>A | VdW |
| | F33 | C A G M O S W<br>D B H L N R V | VdW | | H25 | U<br>F | VdW |
| | | M Q<br>L P | π - π Stack | | E41 | G<br>H | VdW |
| | E41 | G Q W I<br>H P V X | VdW | L10 | P52 | A C S W I<br>B D R V X | VdW |
| | R56 | O P R X B<br>N Q S I A | VdW | L11 | R56 | X H P<br>I G Q | VdW |
| | E40 | S U I O M K<br>R T X N L J | VdW | | H25 | O G S<br>J D N | VdW |
| | G34 | B L N V<br>A M O W | VdW | | P52 | S A<br>R B | VdW |
| | G48 | B R<br>Q K | VdW | | W51 | M O K Q G U W<br>L N J P H T V | VdW |
| | D17 | D J L<br>G O E | VdW | | E41 | K M Q I C G<br>J L P X D H | VdW |

**Table S5.** Residues involved in interchain interactions between the $_6$VPFSLL$_{11}$ motif and NTR residues in the 24-mer structure of human HSPB1. For each amino acid (AA), the table shows the pair of chains involved in the interaction and the interaction type.

| Interchain interactions between the $_{179}$ITIPVTFE$_{186}$ peptide and the lateral grooves of the ACD | | | |
|---|---|---|---|
| **Motif** | **Lateral groove residue** | **Chains in contact** | **Type** |
| I179 | - | - | - |
| T180 | L109 | D H J V X<br>C G K W I | H-bond |
| | | J V<br>K W | VdW |
| I181 | - | - | - |
| P182 | L109 | A K Q<br>B J P | VdW |
| | S155 | A R<br>B S | VdW |
| | L157 | E K U W<br>F J T V | VdW |
| V183 | L155 | N P X<br>O Q I | VdW |
| | L156 | D H N V<br>C G O W | VdW |
| | L157 | A I<br>B X | VdW |
| T184 | L157 | E<br>F | VdW |
| | L157 | Q<br>P | H-bond |
| | S154 | T<br>U | H-bond |
| F185 | S154 | N V<br>O W | VdW |
| | S156<br>S156 | D N P L F<br>C O Q M G | π - H-bond |
| | | D F M P R T V I L H E<br>C E L Q S U W X M G F | VdW |
| | L157 | Q<br>P | H-bond |
| | | E<br>F | VdW |
| E186 | L157 | Q<br>P | H-bond |

**Table S6.** Interchain interactions in the 24-mer structure of human HSPB1 between the $_{179}$ITIPVTFE$_{186}$ peptide and the lateral grooves of the ACD framed by the β4 ($_{109}$LTVKT$_{113}$) and β8 ($_{153}$VSSSL$_{157}$) strands.

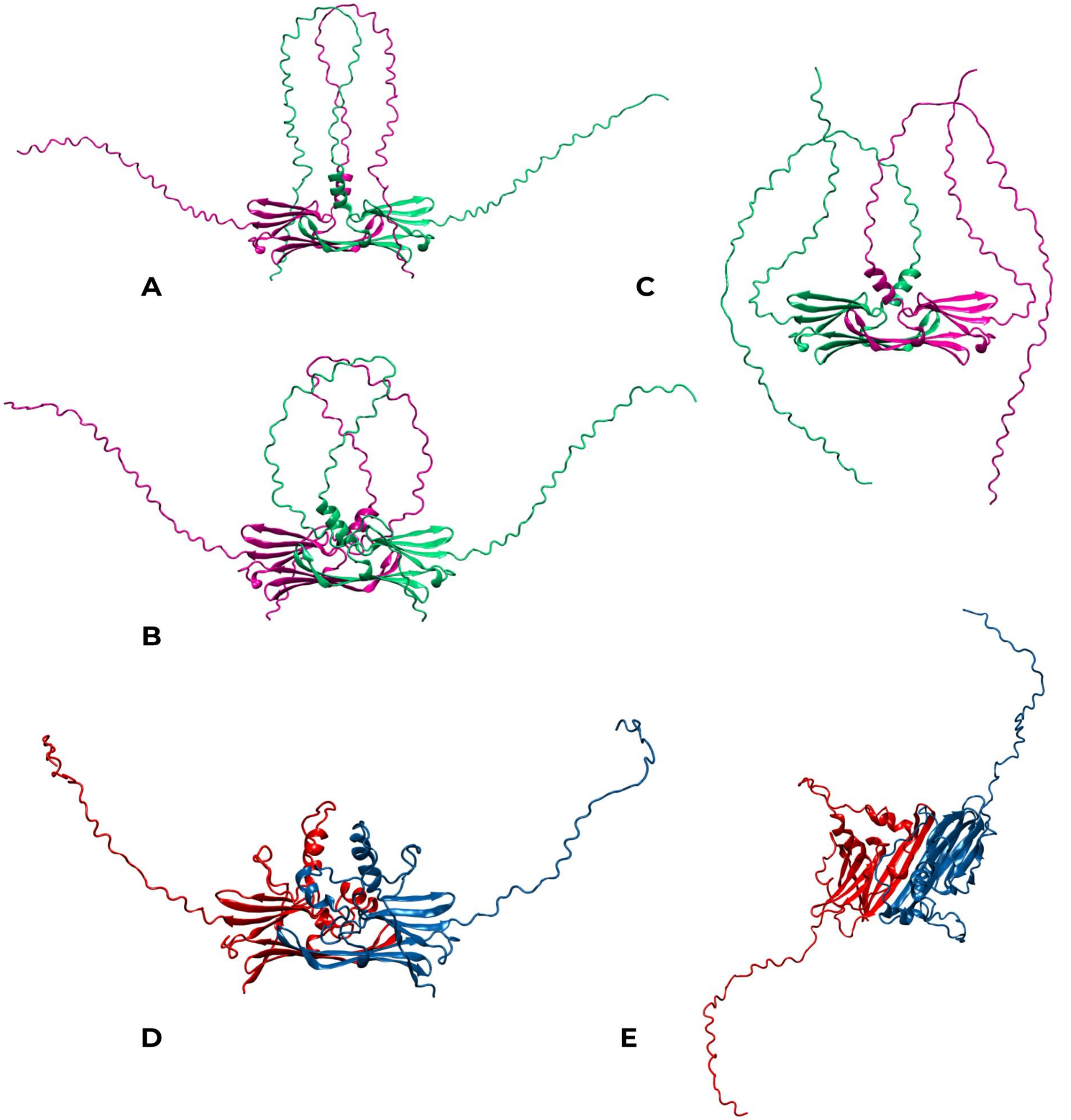


**Figure S2.** Representative conformations of the disordered NTR and CTR in human HSPB1 models generated with AlphaFold. Models A and B depict interchain and intrachain interactions between the NTR and the ACD, with the CTR remaining in solution. Model C shows a conformation where neither the NTR nor the CTR interact with the ACD. The purpose of this modeling was to determine whether interactions between the ${}_{6}$VPFSLL${}_{11}$ motif of the NTR and the ${}_{179}$ITIPV${}_{183}$ motif of the CTR with the ACD could be captured, regardless of the specific conformations adopted by the rest of each disordered region. This approach was necessary because the available structural information for these regions at dimeric level was derived from a phosphomimetic construct rather than from the canonical form of human HSPB1, which was used in this modeling [13]). Models D and E display two perspectives of the same conformation from a dimer model generated using HSPB1's phosphomimetic sequence, modeled with ColabFold [59] and employing PDB 4MJH as a template. In the highest-ranked model among the five provided by the server, the distal (which adopts a β-sheet structure upon binding to the lateral grooves), aromatic, conserved, Trp-rich, inserted, and boundary segments adopt conformations consistent with the experimental descriptions by Clouser et al. for each region, while the CTR remains in solution.

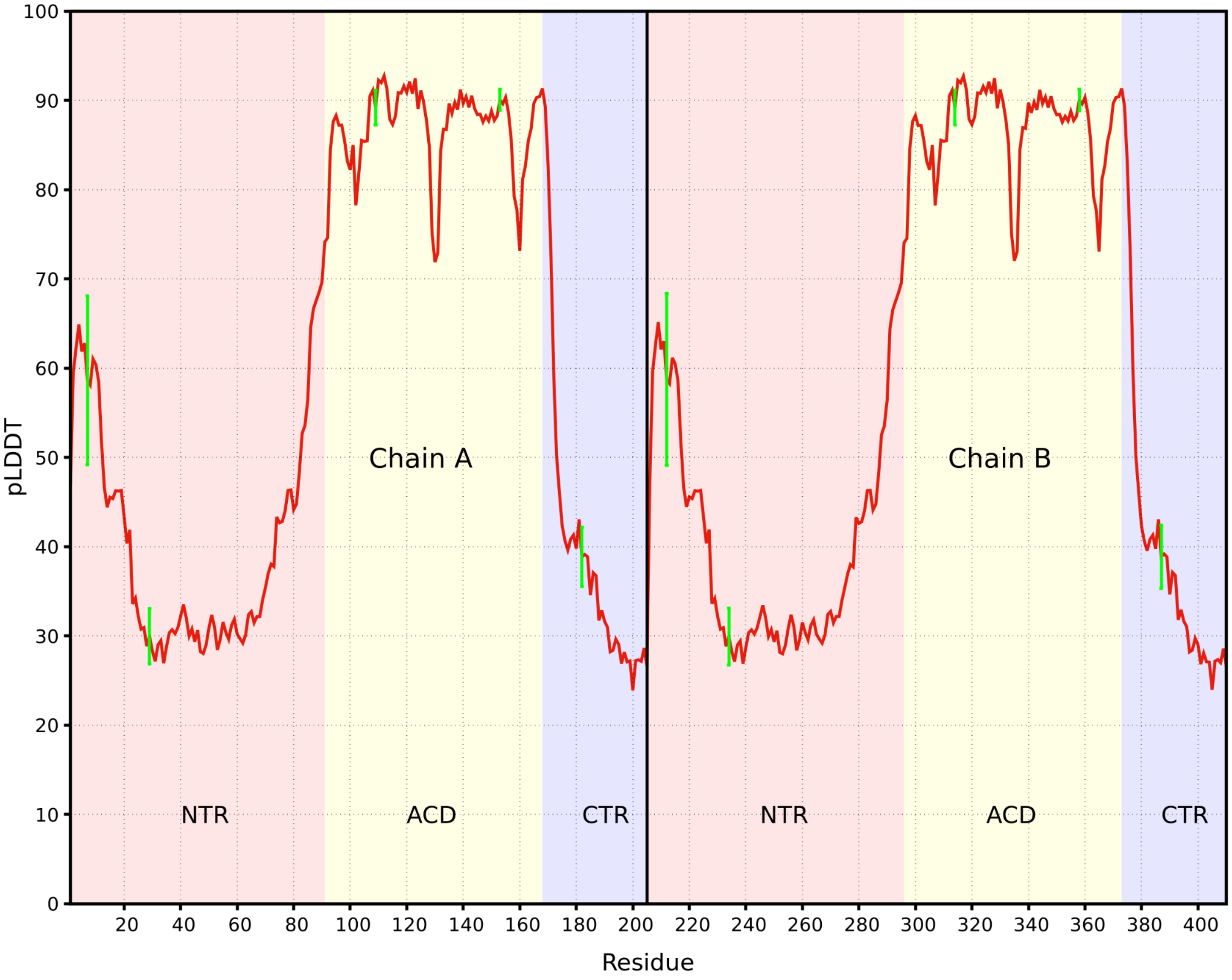


**Figure S3.** Average pLDDT values for each residue across 500 models of the human HSPB1 dimer generated with AlphaFold. Residues 1 to 205 correspond to Chain A, and residues 206 to 410 correspond to Chain B. Background colors highlight the residue ranges for the NTR, ACD, and CTR. Green error bars indicate the standard deviation for key residues across different regions: P7 (from the $_{6}$VPFSLL$_{11}$ motif), F29 (from the conserved $_{26}$SRLFDQXFG$_{34}$ motif), L109 (β4 strand), V153 (β8 strand), and P182 (from the $_{179}$ITIPV$_{183}$ motif).
